# Environmental and genetic disease modifiers of haploinsufficiency of A20

**DOI:** 10.1101/2022.03.19.485004

**Authors:** Nathan W. Zammit, Paul E. Gray, Owen M. Siggs, Jin Yan Yap, Amanda Russell, Daniele Cultrone, Joanna Warren, Stacey N. Walters, Robert Brink, David Zahra, Deborah L. Burnett, Velimir Gayevskiy, Andre E. Minoche, John B. Ziegler, Maria E. Craig, Melanie Wong, Paul Benitez-Aguirre, Juliana Teo, Mark J. Cowley, Marcel E. Dinger, Stuart G. Tangye, Catherine Burke, Tri G. Phan, Christopher C. Goodnow, Shane T. Grey

## Abstract

Monogenic diseases can often manifest diverse clinical phenotypes and cause diagnostic dilemmas. While monoallelic loss-of-function variants in *TNFAIP3* (Haploinsufficiency of A20; HA20) cause a highly penetrant autoinflammatory disease, the variable expressivity suggest a role for additional genetic and environmental disease modifiers. Here, we identify critically ill children who inherited a family-specific *TNFAIP3* deletion from one of their otherwise healthy parents. Each of the probands also inherited *in trans* a subtle loss-of-function I207L *TNFAIP3* variant that is common in Oceania, originally introgressed from Denisovans. Modelling this compound heterozgous state in mice under specific pathogen free conditions demonstrated a reduced threshold to break immune tolerance. Exaggerated immune responses were precipitated by inheriting the two genetic hits on the *TNFAIP3* checkpoint coupled with increasing the microbial challenge to immune tolerance, either by co-housing with pet store mice carrying a wild microbial burden or by transient dietary exposure to a chemical that diminishes the intestinal mucin barrier separating gut microbes from immune sensing systems. These data illuminate second-hit genetic and environmental modifiers contributing to complex inflammatory and autoimmune disease. Increased mechanistic understanding of the presence and contribution of disease modifiers will aid diagnostic and prognostic patient stratification and potentially reveal novel therapeutic opportunities.

## INTRODUCTION

*TNFAIP3* (tumor necrosis factor, alpha-induced protein 3) encodes A20, a highly conserved and evolutionarily ancient protein that serves as a key immune tolerance checkpoint by negatively regulating tissue inflammation and immune responses by inhibiting NF-κB signaling (1-4). A20 inhibits NF-κB signalling though catalytic and non-catalytic mechanisms. A20’s ovarian tumor domain (OTU) and Zinc finger 4 region exert K63-deubiquitinating and K48-ubiqutinating activity, respectively, and co-operate to inactivate key signaling substrate proteins and target them for proteasomal degradation (5-8). In contrast, A20’s Zinc finger 7 binds to linear ubiquitin chains disrupting IKK activation (9, 10). *TNFAIP3* transcription is induced by NF-κB (11-13) and A20’s inhibitory activities are activated by IKKβ-mediated phosphorylation at Ser 381 (9, 14, 15), forming a negative feedback loop on NF-κB signaling.

Common variants at the *TNFAIP3* locus are associated with multiple polygenic autoimmune and inflammatory diseases (16), including type 1 diabetes, rheumatoid arthritis, systemic lupus erythematosus, psoriasis and Crohn’s disease highlighting the importance of *TNFAIP3* in human health and disease. In contrast, haploinsufficiency of A20 (HA20) caused by *TNFAIP3* deletion or protein-truncating variants causes an autosomal dominant, childhood-onset inflammatory disease, reminiscent of Behcet’s disease, with apparent complete penetrance (17). Subsequent findings have reported HA20 in patients with diverse autoimmune and inflammatory dysregulatory conditions including autoimmune lymphoproliferative syndrome (ALPS), systemic lupus erythematosus, type 1 diabetes, hepatitis and interstitial lung disease (18-24).

An emerging feature of HA20 is its association with diverse clinical presentations, phenotypes, and disease severity, even in individuals carrying the same variant, as well as a roughly two times higher incidence in females then men (19, 23-25). The recognition of diverse phenotypes, age of onset and gender bias, suggests a significant role for environmental and or genetic disease modifiers in patients with HA20. Such interactions have been described for Mendelian diseases, including the identification of genes that can modify cystic fibrosis phenotypes in association with mutations in the *CFTR* gene (26), and both gene and environmental factors that influence Huntington’s disease progression (27). However, few have been identified in the context of monogenic immune dysregulatory conditions.

Here, we identify critically ill children who inherited a *TNFAIP3* deletion from one of their otherwise healthy parents, challenging the hypothesis that A20 deficiency alone is always fully penetrant for disease. We now explain these apparently contradictory findings by providing evidence from mouse models for two additional disease modifiers contributing to HA20 in these patients. The first disease modifier is an inherited archaic *TNFAIP3* genetic variant that causes a subtle loss of A20 function, while the second disease modifier came in the form of an environmental challenge that altered microbiome composition. HA20 is a new disease and increased mechanistic understanding of the presence and contribution of disease modifiers will aid diagnostic and prognostic patient stratification and potentially reveal novel therapeutic opportunities.

## RESULTS

Here, we studied two families, Family 1 of mixed Scottish-Fijian/Samoan ancestry and Family 2 of European and New Zealand Māori ancestry (Figure 1A), each with a child with early-onset inflammatory and autoimmune manifestations (Figure 1B, Supplementary Figure 1A,B). Both Child 1 (Family 1) and Child 2 (Family 2) presented with inflammatory bowel disease (IBD), autoantibodies, and cytopenia. Child 1 also exhibited ocular inflammation, subclinical thyroid disease and rhinitis, whereas Child 2 also presented with mouth and genital ulcers, arthralgia/arthritis and type 1 diabetes. Whole genome sequencing of the parent-proband trios revealed each family harbored a unique deletion at the *TNFAIP3* locus, present in both the proband and one unaffected parent respectively. The deletion in Family 1 spanned the entire *TNFAIP3* locus (g.137683250-139345585del; Figure 1C) and was confirmed by CGH array, while Family 2 harboured a 2bp frameshift deletion predicting a premature stop codon truncating the normal protein sequence (p.(N371Sfs*17)) (Figure 1D, Supplementary Figure 2). However, neither of the families had a history of childhood-onset inflammatory disease, despite the fact that an unaffected parent in each pedigree aged 59 and 40 respectively, carried the same heterozygous *TNFAIP3* loss of function allele as the affected proband (Figure 1A). Analysis of human sequence data from people without early onset disease (gnomAD v2.1.1) revealed only 2 from 141,456 individuals with heterozygous protein-truncating *TNFAIP3* mutations, compared to an expected rate of 32 individuals based on coding region size and composition (pLI=1, LOEUF=0.2) (28). Studies in vertebrate model organisms also reveal that HA20 does not always drive a disease phenotype, as mice and zebrafish heterozygous for *Tnfaip3* deletion present as healthy when housed under specific pathogen free (SPF) conditions (1, 3). This suggested that additional, but hitherto unknown, disease modifiers cooperate with *TNFAIP3* haploinsufficiency in these families to drive their disease phenotypes.

**Figure 1:**
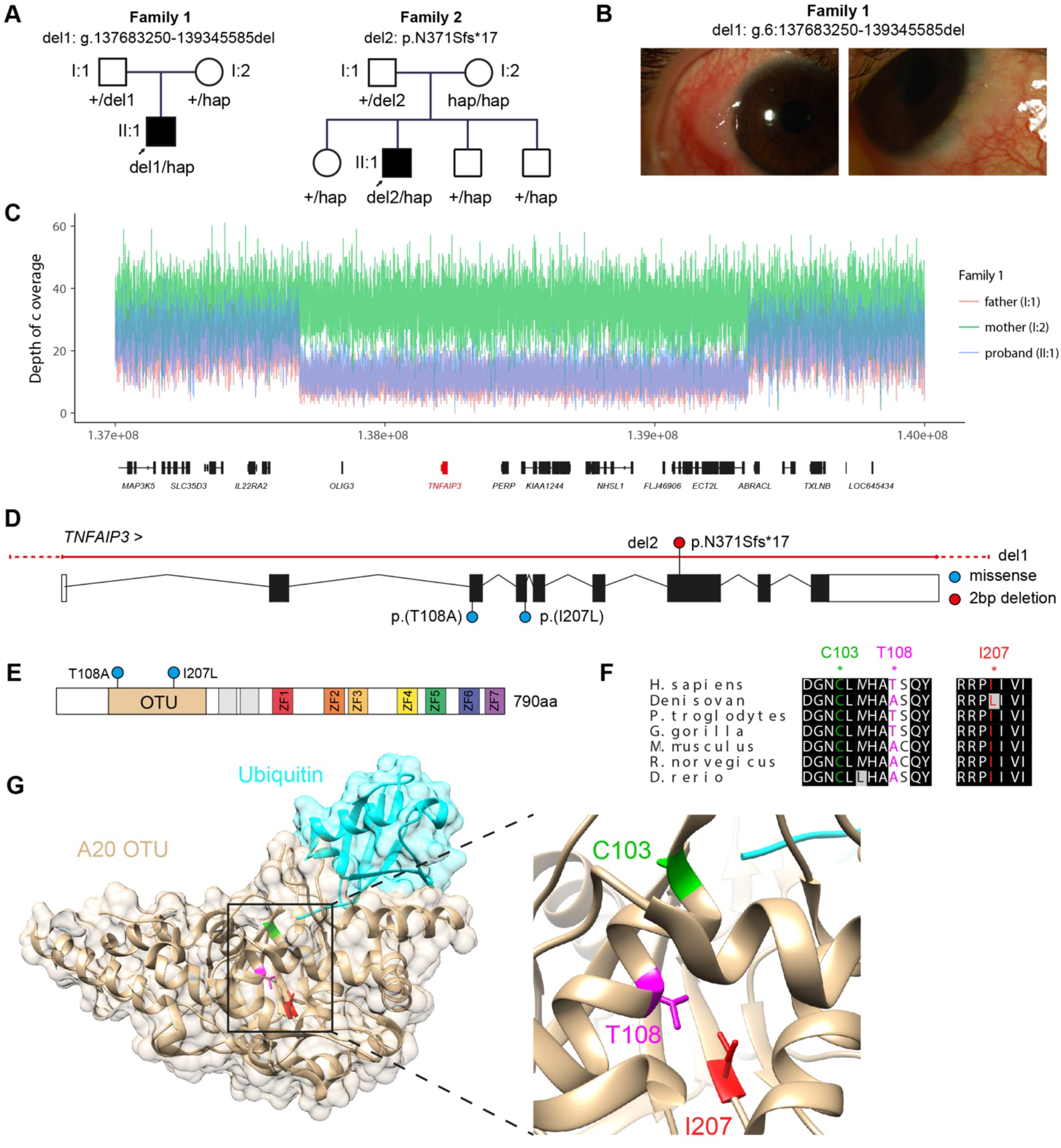
HA20 probands harbour a coding denisovan introgressed *TNFAIP3* allele in *trans* to A20 deletion. **(A)** Family trio’s of identified probands. (**B)** Photo of ocular inflammation from proband of family 1. (**C)** Sequencing coverage map depicting deletion of locus spanning *TNFAIP3. (***D)** Schematic demonstrating the location of HA20 variants del1 from family 1 and del2 from family 2. Each HA20 deletion is located *in trans* to two *in cis* missense variants T108A and I207L, located in exons 3 and 4 of *TNFAIP3*, respectively. T108A and I207L formed part of a larger haplotype adaptively introgressed from Denisovan Hominins (15, 29), referred to as hap in A. (**E)** Location of *TNFAIP3* hap missense variants, T108A and I207L in reference to the primary protein structure of A20. OTU = ovarian tumor domain. ZF = Zinc finger. **(F)** Conservation and (**G)** location of T108 and I207 alleles in context with the OTU domain catalytic 103 (5LRX)(64).

To explore the potential of additional genetic modifiers, we searched for *TNFAIP3* variants *in trans* with each deletion. In both probands we identified the same pair of missense variants *in cis*, p.(T108A) and p.(I207L) (Figure 1D), which formed part of a larger haplotype consisting of multiple coding and non-coding variants (15, 29). Both missense variants are within the OTU domain of A20 (Figure 1E-G), and were rare within public variant databases (gnomAD v2.1.1 allele frequency 0.0001169-0.0001202). We have previously shown, these variants are enriched in populations in Oceania (15), representing 15-45% of all alleles in Polynesia (e.g. Vanuatu), ∼22% of alleles in Martu Australians, 31-48% of *TNFAIP3* alleles in people of the eastern Indonesian islands (e.g. Flores), and 32-100% of alleles in Papuans. Analysis of the genetic history of this haplotype revealed it was inherited by anatomically modern humans from Denisovans ∼50 kya (15, 29). The *TNFAIP3* Denisovan haplotype (hap) contains other non-coding variants found not to impact *TNFAIP3* expression (15). In addition, I207 is predicted to be the functional coding allele, as in contrast to T108, I207L is invariant across most jawed vertebrates (Figure 1F) and has a higher *in-silico* deleterious score (CADD score 23.2; (15)). In transfected cells expressing *TNFAIP3* cDNA, the T108A;I207L substitutions decreased A20 phosphorylation by IKKβ to 80% of wild-type, creating a subtle loss of NF-κB inhibitory function (15). In both families, the T108A;I207L Denisovan haplotype was inherited from one healthy parent and the *TNFAIP3* deletion from the other healthy parent (Figure 1A). Hence, it was conceivable that co-inheritance of *TNFAIP3* haploinsufficiency and a subtle loss of function Denisovan allele explained disease only in the child with both genetic hits in compound heterozygous configuration.

To test if coinheritance of the two alleles had additive biochemical affects we performed Western blot analysis of peripheral blood mononuclear cells (PBMCs) from healthy donors, the heterozygous parents and the affected patients. A20 protein levels were decreased in each of the deletion-carrying parents compared to healthy donors (Hd) (Figure 2A, B & Supplementary Figure 3. A, C). Neither *TNFAIP3* deletion 1 (+/del1), spanning the entire *TNFAIP3* locus, or deletion 2, a frameshift deletion p.(N371Sfs*17), encoded proteins that could be detected by the antibody probe raised against the C-terminus region (aa 440-790) of A20. Indeed, mRNA analysis of *TNFAIP3* levels using probes spanning exons 2 and 4 show a dose dependent reduction for individuals in family 1 harboring a heterozygous deletion of the entire *TNFAIP3* locus compared to 15 healthy donors (Figure 1C). Heterozygous carriers of the I207L allele (+/hap) exhibited similar A20 protein levels to healthy donors, with probands (hap/del) exhibiting levels similar to their +/del parent (Figure 2A, B & Supplementary Figure 3A, C). Active IKKβ also phosphorylates another NF-κB inhibitory protein, IκBα, which then undergoes K48-linked ubiquitination and degradation to release active NF-κB (30, 31). Western blot analysis revealed levels of IκBα were greatly decreased in PBMCs from the del/hap probands in both families even under non-stimulated conditions (Figure 2A, B; Supplementary Figure 3B, D). For family 1, PBMCs from the healthy parent with the *TNfAIP3* deletion (+/del) exhibited more rapid loss of IκBα following TNF stimulation compared to PBMCs from healthy donors (Figure 2A; Sup. Figure 3B), as previously reported for HA20 (17). However, this effect was less than the del/hap proband from this family (Figure 2A; Sup. Figure 3B). In family 2, the healthy +/del parent exhibited greatly decreased IκBα similar to the del/hap proband (Figure 2B; Sup. Figure 3D).

**Figure 2:**
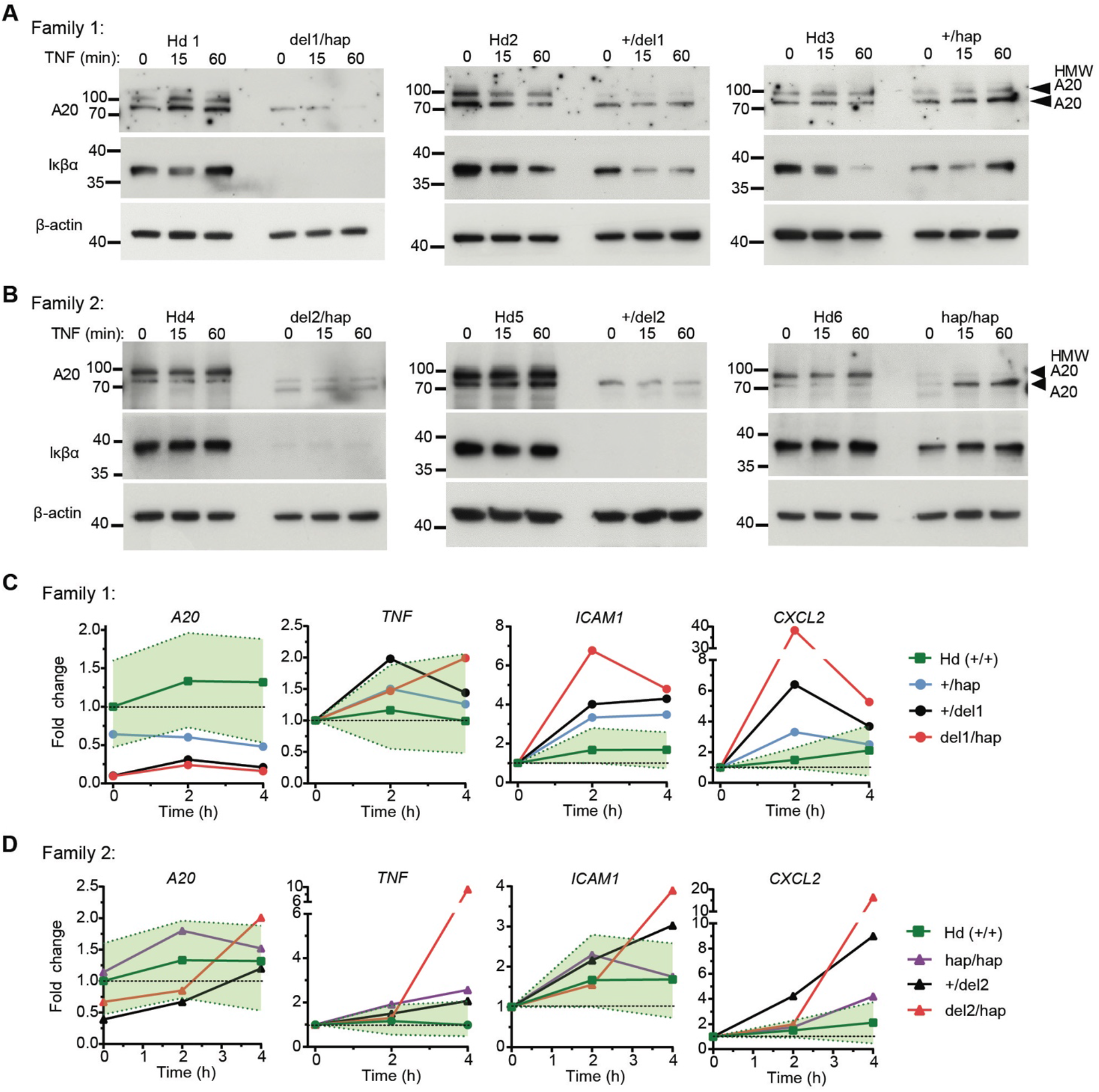
Analysis of peripheral blood mononuclear cells (PBMCs) from parent-proband Trios. **(A-B)** Immunoblot analysis of PBMCs from parent-proband trios from family 1 **(A)** and family 2 **(B)**. Probands from each family harbor a *TNFAIP3* deletion (del1 or 2) and the *TNFAIP3* T108A;I207L oceanic haplotype (hap) in *trans* (del/hap). Parents are either heterozygous for a *TNFAIP3* del or heterozygous or homozygous for the *TNFAIP3* hap allele. PBMCs were left untreated or stimulated with TNF for 15 or 60 min. Proteins assessed included A20, IκBα and β-actin (loading control). A20 high-molecular weight bands (HMW) representing phospho-A20 are indicated (9, 14, 15). Healthy donors = Hd. Immunoblots of PBMC lysates from +/Hap and hap/hap parents (right panels) have been previously reported (15)(ref). **(C-D)**, RT–PCR analysis of *TNFAIP3, TNF, ICAM1* and *CXCL2* gene expression in PBMCs from proband trios in A and B. Probands (red), +/hap (blue), hap/hap (purple), +/del (black) and 15 healthy donors (green) +/- range (shaded light green).

RT-PCR analysis of PBMCs was used to measure mRNA encoding NF-κB induced inflammatory genes (30) before and following TNF stimulation. These target genes were more strongly induced in cells from del/hap probands than in PBMCs from *TNFAIP3*^+/+^ donors. Cells from +/del individuals exhibited an intermediate phenotype, while +/hap cells showed a subtle hyper-inflammatory response (Figure 2C, D). Collectively, these data are consistent with an additive biochemical effect of coinheriting a *TNFAIP3* null/truncating and *TNFAIP3* hypomorphic allele, but are not conclusive because of the small number of donors with the relevant genotypes.

Given the limited availability of matched +/del and hap/del individuals to make conclusions about additive effects, we generated two mouse lines by Cas9/CRISPR gene editing on the genetically homogeneous C57BL/6 background, transmitting either a *Tnfaip3* truncating mutation (p.Ala240Valfs*7, referred to as del) or a *Tnfaip3* T108A;I207L allele (hap) (15) (Supplementary Figure 4A, B). Next we crossed both lines so that offspring would be wild-type, heterozygous truncating (+/del), heterozygous T108A;I207L (+/hap), or compound heterozygous (hap/del) to model the human family members (Figure 3A). *Tnfaip3* +/hap, +/del and hap/del mice reared in SPF conditions exhibited normal weights at weaning and at 8 weeks of age (Figure 3B; Supplementary Figure 4C, D). Heterozygous loss of full length A20 due to the truncating allele was confirmed by a decrease to 50% of wild-type A20 protein levels in isolated +/del and hap/del thymocytes (Figure 3C, D). Western blot analysis of TNF-stimulated thymocytes revealed decreased IκBα protein levels even under non-stimulated conditions for compound heterozygous hap/del cells that was not observed for thymocytes from +/hap or +/del thymocytes (Figure 3C, E). *Tnfaip3* +/del cells exhibited a trend to increased stimulus-dependent loss of IκBα compared to +/hap and WT cells (Figure 3C, E). These findings highlight an additive role for +/del and +/hap *Tnfaip3* alleles in restraining NF-ĸB inhibitory activity.

**Figure 3:**
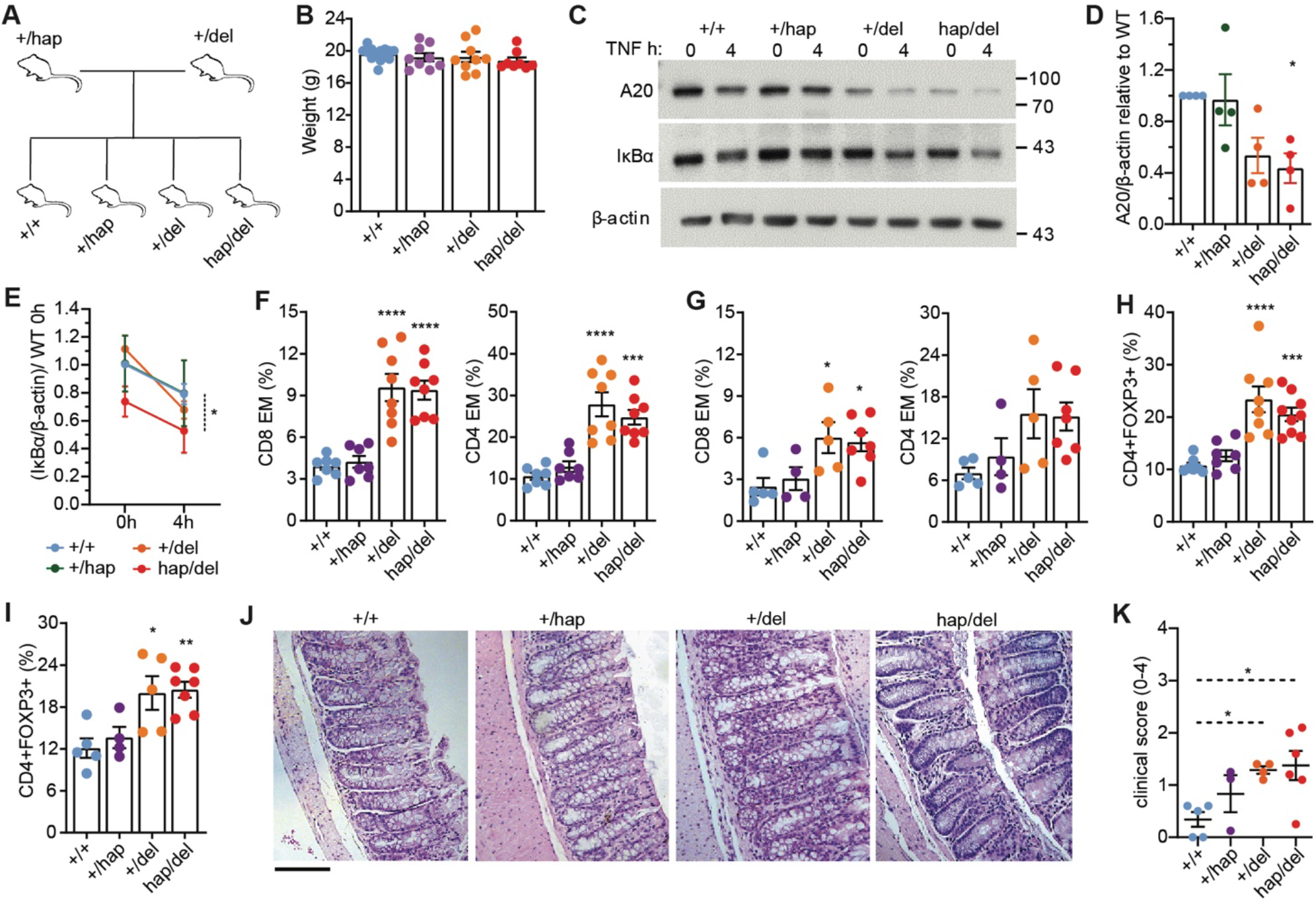
Molecular and phenotypical analysis of hap/del mice. **(A)** Pedigree tree illustrating breeding strategy for *Tnfaip3* hap/del line. **(B)** Weight of 8-week-old female *Tnfaip3* hap/del mice of the indicated genotype. **(C)** Representative immunoblot of thymocyte lysates isolated from mice with indicated genotypes and stimulated with TNF for indicated times. Proteins probed include A20, IκBα and loading control β-actin. Cumulative densitometry is shown for **(D)** A20 and **(E)** IκBα with each point representing thymocytes from an individual mouse. **(F-G)** Percentage CD44hiCD62Lo effector memory cells among CD8+ or CD4+ T cells from the spleen (F) or Mesenteric lymph node (G) of mice with the indicated genotype. **(H-I)** Percentage of FOXP3+ regulatory T cells among CD4+ cells from the spleens (H) or mesenteric lymph nodes **(I)** of mice of the indicated genotypes. Flow cytometric data are cumulative from 3 independent experiments. **(J)** Representative sections showing H&E-stained colons from female mice of the indicated genotype. Scale bar = 200 µm. **(K)** Clinical score of representative colons shown in (J). Statistical analysis was performed by one-way anova ANOVA with Tukey post hoc test, and data shown as the mean +/- s.e.m: *P<0.05; \*\**P <* 0.01; \*\*\**P <* 0.001; \*\*\*\**P <* 0.0001.

In contrast to the exaggerated biochemical response in the compound heterozygous mice, flow cytometric analysis revealed comparable increases in the proportions of CD44^hi^CD62^low^ effector memory CD8 and CD4 T cell subsets in the spleen (Figure 3F) and mesenteric lymph nodes (MLN, Figure 3G) of *Tnfaip3* +/del and hap/del mice compared to +/hap and wild-type littermates (Supplementary Figure 5A-F). *Tnfaip3* +/del and hap/del mice also exhibited a comparably increased proportion of FOXP3+ Tregs among CD4 T cells (Figure 3H, I), and increased number of B cells in the spleen (Supplementary Figure 5G). However, only hap/del mice exhibited significantly increased numbers of germinal center B cells in the spleen and MLN under SPF conditions (Supplementary Figure 5I, J).

A major clinical phenotype in HA20 patients included gut pathology (Supplementary Figure 1A). Interestingly, mucosal inflammation is a unifying symptom of human HA20 disease (24, 32) and spontaneous gut inflammation observed in *Tnfaip3* gene-targeted mice is driven by exaggerated responses to gut microbiota (33). Analysis of colon sections from SPF *Tnfaip3* hap/del mice, revealed immune infiltrates with a mild clinical score similar to *Tnfaip3* +/del mice, which was higher than that observed in *Tnfaip3* +/hap or wild-type littermates (Figure 3J, K).

The results above indicated that *Tnfaip3* haploinsufficiency (+/del) in mice causes subclinical immune dysregulation that is not markedly increased by co-inheriting the hypomorphic I207L allele when the microbial flora is highly controlled in an SPF facility. To better replicate natural microbial stimulation of the immune system, we co-housed the *Tnfaip3* mutant mice with “dirty” mice harboring a diverse microbiome. Cohousing mice with non-laboratory wild mice has been shown to facilitate the horizontal transfer of microbiota, maturing the immune system (34, 35). Deep sequencing of bacterial 16S rRNA of fecal pellets from the co-housed mice showed horizontal transfer resulting in a more diverse gut microbiota compared to their SPF counterparts, consisting of an increase in the relative abundance of Prevotella, Odoribacter, Helicobacter, Muscispirillum and a decrease in Akkermansia. However, no discernible difference in principle microbial components was observed among co-housed *Tnfaip3* hap/del, +/del, +/hap and +/+ animals (Figure 4A; Supplementary Figure 6A, B). Cohousing did not adversely affect the weight or survival of *Tnfaip3* wild-type, +/hap, +/del or hap/del mice (Supplementary Figure 6C), but increased the percentage of CD44^hi^ effector/memory CD4 and CD8 T cells in the blood within 1 week of cohousing (Supplementary Figure 6D, E). Following 8 weeks of cohousing, clinical scores for colon inflammation tended to be higher in *Tnfaip3* hap/del mice than +/del animals, with marked mononuclear infiltrates and goblet cell hyperplasia (Figure 4B, C). In the MLN draining the gut, the proportion of effector/memory cells among CD8 and CD4 T cells was increased in +/del mice and further increased in hap/del compound heterozygotes (Figure 4D, E). A similar finding was observed when measuring total CD44+ CD8 T cells (Supplementary Figure 5D), and CD4 central memory T cells in MLNs (Supplementary Figure 5E, F). Cohousing with dirty mice did not segregate Treg frequencies between *Tnfaip3* +/del and hap/del genotypes (Figure 4F, G) but significantly decreased mean FOXP3 protein levels per cell in splenic Tregs of *Tnfaip3* hap/del mice compared to +/del mice (Figure 4H). In cohoused mice, percentage of splenic germinal center B cells was progressively increased according to *Tnfaip3* genotype consistent with additive effects of the del and hap mutations (Supplementary Figure 5 H-J).

**Figure 4:**
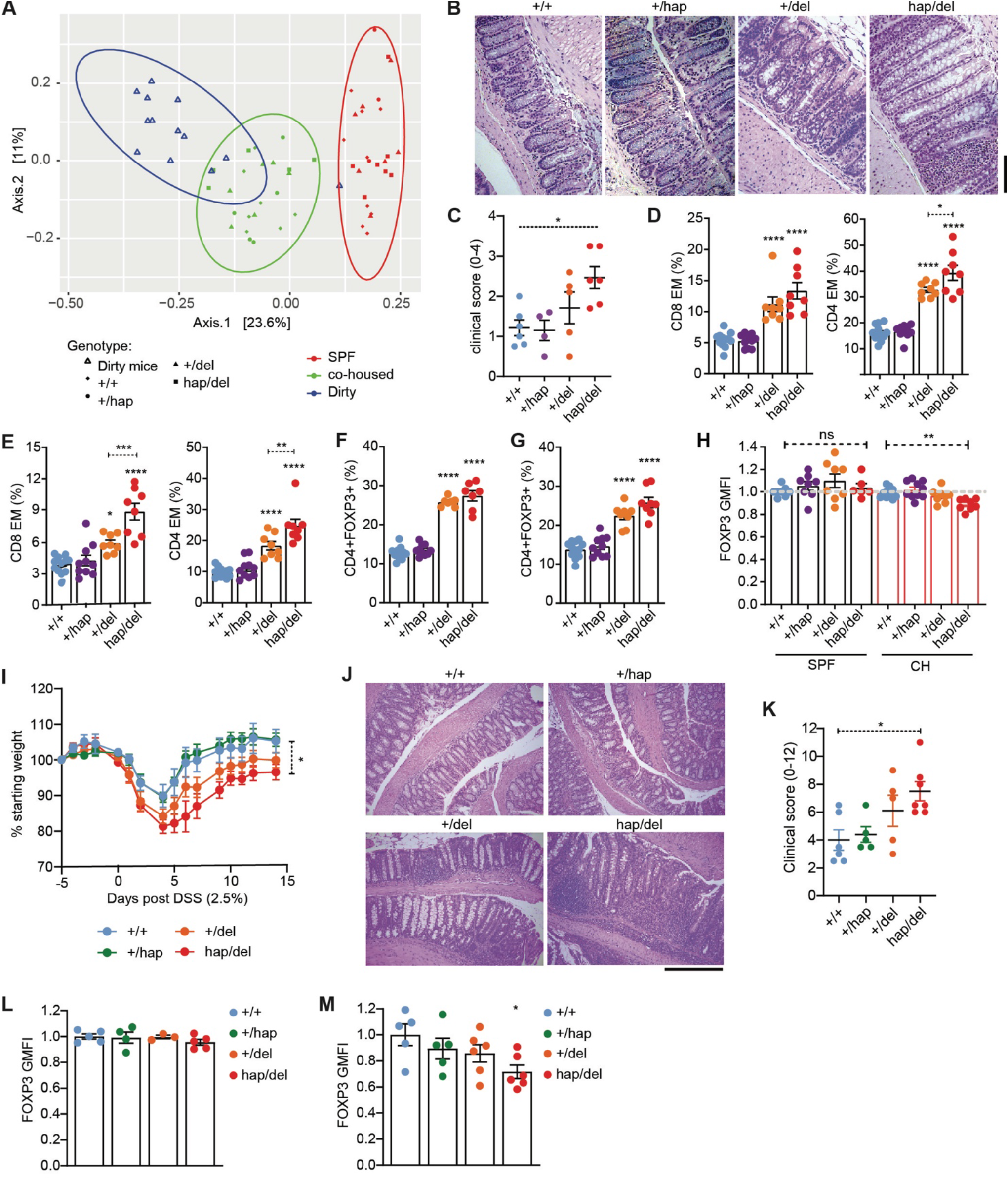
Microbiome and DSS challenge uncovers differential effects for hap/del mice compared to +/del alone. **(A)** Unweighted unifrac PCoA showing cohoused *Tnfaip3* hap/del mice take on microbiota from dirty mice (P<0.001, PERMANOVA dirty mice vs SPF vs co-housed). Genotypes indicated by symbols. **(B)** Representative sections showing H&E-stained colons from female *Tnfaip3* hap/del mice 8 weeks post cohousing with dirty mice (20x magnification; scale bar = 200 µm), and **(C)** cumulative clinical score. **(D-E)** Percentage of CD44hiCD62Lo effector memory CD8+ or CD4+ T cells from the spleen (**D**) or mesenteric lymph node (**E**). **(F-G)** Percentage of CD4+FOXP3+ regulatory T cells from the spleens (**F**) or mesenteric lymph nodes (**G**) of mice of the indicated genotypes. **(H)** FOXP3 GMFI for CD4+FOXP3+ splenic Tregs without cohousing (black bars) and post 8 weeks cohousing (red bars). FOXP3 GMFI normalised to the average GMFI of +/+ samples **(I)** Percent change in starting weights of mice with indicated genotypes given 2.5% DSS in drinking water for 5 days. Data is cumulative of two independent experiments, +/+ n=6; +/hap n=5; +/del n=8; hap/del n=8. Statistical significance determined by area under the curve analysis. **(J)** Representative haematoxylin and eosin-stained colon sections 14 days post DSS treatment and **(K)** cumulative clinical scores. **(L)** FOXP3 GMFI for CD4+FOXP3+ splenic Tregs under SPF conditions and **(M)** 14 days post DSS treatment. FOXP3 GMFI normalized to the average GMFI of +/+ samples. Statistical analysis was performed by one-way ANOVA with Tukey post hoc test unless otherwise stated, and data shown as the mean +/- s.e.m: \**P <* 0.05;\*\*\*\**P <* 0.0001. ns = not significant.

To test a different environmental stressor, we next challenged *Tnfaip3* hap/del mice maintained in SPF conditions by adding 2.5% dextran sodium sulfate (DSS) to drinking water. DSS thins the intestinal mucin barrier that normally promotes bacterial tolerance by physically separating most gut microbiota from the intestinal epithelium. DSS was given for 5 days, then removed (day 0) and weight loss and recovery monitored for 2 weeks. Weight loss was exaggerated and recovery delayed in *Tnfaip3* +/del animals and these effects were worsened in hap/del compound heterozygous animals (Figure 4I). Only *Tnfaip3* hap/del mice exhibited significant weight loss over the test period compared to wild-type littermates. Two weeks post-DSS removal, colons were collected and assessed for inflammatory cell infiltrate, erosion and ulceration, hyperplasia and submucosa oedema and goblet cell and crypt loss. *Tnfaip3* hap/del mice exhibited the highest clinical scores (Figure 4J, K). DSS challenge caused an increase in effector/memory CD44^hi^ CD8 T cells and FOXP3^+^ Tregs for all genotypes assessed (Supplementary Figure 7A-D). In Tregs, *Tnfaip3* hap/del mice treated with DSS exhibited a significant decrease in mean FOXP3 protein per cell compared to wild-type littermates treated with DSS, while +/del animals tended to be intermediate (Figure 4L, M). DSS treatment also increased the frequency of neutrophils in the spleen and MLNs, with *Tnfaip3* hap/del mice exhibiting higher frequency of splenic neutrophils compared to +/del alone and wild-type littermates (Supplementary Figure 7E, F).

## DISCUSSION

The findings here reveal how rare and common inherited variants in an important immune tolerance checkpoint interact with environmental stressors to precipitate autoimmune and inflammatory disease. Rare deletions in *TNFAIP3* have been considered to cause a fully penetrant autosomal dominant, childhood-onset inflammatory/autoimmune disease by haploinsufficiency (17). However, HA20 patients present with diverse clinical characteristics and variable age of onset suggesting a role for additional disease modifiers. Here we describe two children with clinical HA20 syndrome who inherited a family-specific, heterozygous inactivating *TNFAIP3* deletion from one well parent and a subtle loss-of-function *TNFAIP3* allele that is common in Oceania, from their other healthy parent (15, 29). Ex-vivo analysis of patient cells demonstrates increased inflammation and NF-κB signaling in asymptomatic deletion carriers, which is potentiated in symptomatic probands. This potentiation was replicated in mice rendered compound heterozygous for a deletion and the common Oceanian allele, precipitated by exposing the mice to diverse gut microbes or damaging the mucin barrier between gut microbes and the body’s microbe response systems. The families described here demonstrate that contrary to current consensus (18-22), *TNFAIP3* haploinsuffucuency does not always cause a fully-penetrant childhood-onset inflammatory disease, and that *TNFAIP3*-associated inflammatory disease exemplifies interaction between environmental triggers and common and rare inherited alleles.

Here we show for the first time that some people carrying a heterozygous TNFAIP3 deletion/truncating allele do not develop inflammatory/autoimmune disease. Why does this occur? The animal experiments here support two explanations that are not mutually exclusive. First, asymptomatic carriers may lack second genetic hits – single or multiple common polymorphisms - that further compromise the *TNFAIP3* immune tolerance checkpoint. The ‘gene dose’ effect observed for *TNFAIP3* variants here may be analogous to the potentiation of rare deletions in RBM8A by common *in trans* hypomorphic SNPs in the causation of the thrombocytopenia with absent radii (TAR) syndrome (36). By contributing to compound heterozygous inheritance of HA20, the Denisovan-derived I207L (*Tnfaip3* hap) is a rare example of a variant with relevance both at a population and Mendelian level, analogous to protection from severe Falciparum Malaria afforded by heterozygosity for sickle cell variants (37).

Second, these genetically predisposed, asymptomatic individuals may not have had as much pressure applied to the checkpoint by environmental microbes. These concepts build upon Archibald Garrod’s prototypic recessive “inborn error of metabolism”, alkaptonuria (AKU) (38), and the more common recessive disease phenylketonuria (PKU) (39), where the genetic dose of mutations inactivating enzymes for phenylalanine and tyrosine metabolism cooperates with the dietary load of these amino acids. Above a certain threshold, over time, accumulating metabolic intermediates form toxic polymers on collagen (alkoptonuria) or unmetabolized phenylalanine blocks transporters at the blood-brain barrier leading to developmental delay. In the case of PKU, understanding the gene/environment cooperation enables disease prevention by limiting dietary phenylalanine. For people with an underlying gene deficit in the *TNFAIP3* immune tolerance checkpoint, it will be important to recognise these individuals early and identify ways to minimise microbial stimuli that take them over a threshold into inflammatory or autoimmune disease.

Exposing *Tnfaip3* hap/del mice to ‘dirty mice’ with ‘wild’ microbial flora or transiently introducing DSS in the drinking water (40), revealed a significantly worse outcome for *Tnfaip3* hap/del mice compared to single variant mice. A thick inner mucin layer secreted by intestinal goblet cells normally serves a key role in tolerating microbial flora in the gut lumen. This is achieved by physically separating them from bacterial-sensing receptors on and within intestinal epithelial cells and lamina propria macrophages, notably Toll-like receptors (TLR) and NOD-like receptors (NLR) that activate NF-ĸB. Oral DSS rapidly disrupts the mucin barrier to increase bacterial contact with the epithelium (41, 42). The resulting stimulation of TLR- and NLR-induced NF-ĸB activation will increase pressure on the *TNFAIP3* checkpoint to dampen NF-ĸB signaling and maintain tolerance to the bacteria, providing an explanation for the additive effects of compound heterozygous hap/del *TNFAIP3* mutations on DSS pathology. Consistent with this interpretation, complete deletion of both TNFAIP3 alleles selectively in murine intestinal epithelial cells cooperates with DSS treatment and gut flora to cause a similar inflammatory bowel disease (43).

As opposed to dietary compromise to the mucin barrier, a number of the bacterial species that were expanded in the intestine of the co-housed mice, have specialized mechanisms to penetrate the mucin barrier and adhere directly to intestinal epithelium. Gram-negative Prevotella species of the Bacteroidetes phylum and segmented filamentous bacteria predispose to autoimmune disease (44, 45) by mechanisms that depend upon adherence to intestinal epithelium (46, 47). Mucispirillum colonizes the mucin layer to provoke inflammatory bowel disease in mice lacking NOD2 and CYBB NADPH oxidase (48). Helicobacter hepaticus provokes colitis in IL-10 deficient mice (49). These expansions in bacterial species coincided with a loss of Akkermansia that exhibit anti-inflammatory properties that maintain the mucin barrier (50, 51). In mice with specific pathogen free flora, homozygous loss of TNFAIP3 in intestinal epithelial cells and in intestinal macrophages results in a vicious cycle of microbe-induced overproduction of TNF by macrophages and intestinal epithelial cell hyperproliferation and apoptosis, depleting goblet cells and the mucin barrier (52). Collectively, these results illustrate how genetics, diet and gut microbial flora cooperate to exceed tolerance thresholds.

A significant loss of FOXP3 MFI was observed in *Tnfaip3* hap/del Tregs following cohousing and DSS treatment that was not observed for single variant mice, potentially explaining their segregating phenotypes. Treg activation and expansion is NF-κB dependent via the TCR-CARD11 pathway (53-55). This is further evidenced by expanded Treg populations observed in the *Tnfaip3* variant mice (15). However, chronic inflammation and persistent NF-κB activation can drive FOXP3 instability (56, 57) by preventing FOXP3 hypomethylation, which requires CD28/NF-κB deprivation in developing Tregs (57). These data suggest that the biallelic addition of +/hap and +/del, potentiates NF-κB activation under inflammatory conditions, down-regulating FOXP3 expression. Indeed western blot analysis of TNF stimulated thymocytes show an additive effect for *Tnfaip3* +/hap and +/del variants on the NF-κB inhibitor, IκBα. As A20 haploinsufficiency lowers immune stimulatory thresholds, a loss of FOXP3 expression would likely be detrimental contributing to clinical inflammatory conditions (58-61).

The experimental approaches deployed here possess real potential in the study of both HA20 and polygenic inflammatory bowel disease (IBD). The failure of mouse avatars representing the genetics of human monogenic inflammatory disease to express a clinical phenotype unless specifically challenged is not unique, with *Lrba* gene-deficient animals requiring DSS stimulation to develop clinical disease (62). The models of colitis presented here associated with mice having a more ‘wild-type’ intestinal flora contributes to growing knowledge for human IBD where the search for pathogenic mechanisms is increasingly focused on changes to the intestinal microbiota (63).

In summary, we demonstrate for the first time, incomplete penetrance associated with deletions in *TNFAIP3*. The Oceanic allele that contains the functional p.I207L allele represents a unique example of a common variant contributing to a Mendelian inflammatory disease, and has implications for genomic filtering strategies, which might remove such variants. Furthermore, *TNFAIP3* T108A;I207L which was adaptively introgressed into Oceanian populations and can potentially provide resistance to infection (15), may represent the first example of an immune variant with a population protective benefit, that is Mendelian deleterious. It remains to be seen whether this same variant influences immunity and inflammation in other non-Mendelian diseases in the large number of individuals who carry it today. Finally, our data highlight the significance of ancestry in rare disease penetrance, and the value of studying inherited disease in non-European populations, which are poorly represented in current clinical and population genetic studies.

## AUTHOR CONTRIBUTIONS

CCG – project initiation, coordination, and data interpretation.

PEG, OMS, AR, JBZ, MEC, PB-A, JT, MW, MJC, VG, AEM, MED, CCG – Clinical recruitment, whole genome sequencing analysis of HA20 patients.

NWZ, JYY, SGT, AR, STG – Molecular and immune phenotyping of human PBMCs.

NWZ, DC, STG – molecular characterization of A20 variants.

NWZ, JW, SNW, DC, RB, DZ, DLB, STG – mouse avatar characterization, pathology and disease models.

NWZ, CB, TP, STG – Microbiome challenge (DSS model; cohousing model) and microbiome analysis.

NWZ, PEG, OMS, CCG, STG - analysed data and co-wrote the manuscript.

## ACKNOWLEDGMENTS

We thank the Biological Testing Facility at the Garvan Institute of Medical Research for animal care.

We thank the Clinical Immunogenomics Research Consortium of Australasia for collecting and co-ordinating human samples (CIRCA). https://www.garvan.org.au/research/collaborative-programs/circa/about-circa.

## DATA AVAILABILITY

The data that support the findings of this study are available from the corresponding author upon request

## Methods

### Human subjects

Patients and healthy family members were recruited via the Clinical Immunogenomics Research Consortium Australia (CIRCA) and The Children’s Hospital at Westmead, providing written informed consent under protocols approved by the Sydney *Children’s Hospitals* Network Human Research *Ethics* Committee (SCHN HREC). Blood samples from healthy volunteers were obtained from the Australian Red Cross Blood Service.

### Genome sequencing

Parent-proband trio genomes were sequenced on the Illumina HiSeq X platform using DNA isolated from whole blood, to 30x genome-wide coverage. Libraries were generated using either Illumina TrueSeq PCR-free or TrueSeq Nano. Alignment, variant calling, and annotation was performed as described previously (65-67).

### Mice

To generate *Tnfaip3*^*246tr*^ and *Tnfaip3*^*I207L*^ strains, respective guide RNAs (5ʹ-GGGATATCTGTAACACTCC-3ʹ and 5ʹ-TGACAATGATGGGTCTTCTGAGG-3ʹ) were microinjected into C57BL/6 zygotes in combination with Cas9 mRNA. For *Tnfaip3*^*I207L*^ strain an oligo template was also co-injected (5ʹCTCCTCAGAGCTGAAACTCACCCAGGGAACCTAGAAACTCTCTGAGGCACCTCACCTGAAATGACAATGAG GGGTCTTCTGAGAATGTTGCTGAGGACAAATATGTGGATTTCTTCCAGGGAATTGTACTGAAGTCCACTTCGG GCTGCA-3ʹ). Founder mice carrying the appropriate substitutions were then crossed to C57BL/6, and heterozygous offspring intercrossed to generate homozygous, heterozygous and wild-type littermates. Mice were housed at the Australian Phenomics Facility (Australian National University, Canberra, Australia), or at the Australian BioResources Centre (ABR) (Moss Vale, NSW, Australia).

Animal studies were approved by the Garvan/St Vincent’s or the Australian National University Animal Ethics Committees. All procedures performed complied with the Australian Code of Practice for Care and Use of Animals for Scientific Purposes.

### Microbiome 16S rRNA sequencing and analysis

Fecal pellets from individual mice were stored at −80 prior to processing. DNA was extracted using the PowerFecal DNA extraction kit (MoBio), including blanks as negative controls. 16S rRNA gene libraries were prepared by amplifying the V4 region with primers that contained the 515F (GTGYCAGCMGCCGCGGTAA) and 806R (GGACTACNVGGGTWTCTAAT) sequences, along with 0-3 bp phasers and partial Truseq Illumina adaptors, and KAPA HiFi Hotstart ready mix for 20 amplification cycles. PCR products were diluted 1:200 and amplified for a further 10 cycles with primers that included the TruSeq Illumina adaptors and 6bp sample barcodes on each primer. Both DNA extraction blanks and PCR negative controls were included. Final PCR products were cleaned with 0.8V JetSeq Clean Beads (Bioline) and PCR yield measured with a Quant-it PicoGreen ds DNA assay (ThermoFisher). Samples with a detectable PCR yield were pooled at an equimolar concentration, while negative controls or samples that had no detectable yield were added to the pool at the median volume used for all other samples. The pool was quantified with a Bioanalyser HS DNA chip. The library was sequenced on an Illumina MiSeq using a V2 flow cell with 2 × 250 bp paired end reads.

Sequencing data was assessed for quality using FastQC, and processed using Qiime2 v 2019.10 (68). In brief, primers and spacers were trimmed using cutadapt (69), forward and reverse reads were merged using vsearch (70) and sequences were quality filtered and denoised to amplicon sequence variants (ASVs) using Deblur (71). ASVs were classified using a naive Bayes machine-learning classifier (72) trained on the V4 region of the Greengenes 16S rRNA gene operational taxonomic unit (OTU) database clustered at 99% (accessed from https://docs.qiime2.org/2019.10/data-resources/ in November 2019). A rooted phylogenetic tree was constructed using the phylogeny plu-in and align-to-tree-mafft-fasttree command. Differential relative abundance analysis was performed using the composition plug-in and ANCOM (73).

Further analysis was performed in the R environment. Phyloseq (74) was used to import and transform ASV counts and sample metadata. Samples with less than 10x the largest sequence count of any negative controls were removed from the analysis. ASV counts were log transformed prior to calculating weighted and unweighted unifrac distances using a Phyloseq wrapper of FastUnifrac (75), and used as input for Principal Coordinates Analysis in Phyloseq. Significant differences in clustering were assessed with PERMANOVA (76) via the adonis function of Vegan (77). Faith’s phylogenetic diversity (78) was calculated with Picante (79) and differences in diversity between groups assessed with ANOVA in base R. Data frames were manipulated using dplyr (80) and plots generated with ggplot2 (80).

### Immunohistochemistry

Tissues were fixed in 10% neutral buffered formalin (Sigma-Aldrich), paraffin embedded and sections (5 µm) prepared and stained with hematoxylin and eosin (H&E; Sigma-Aldrich). For clinical scoring 4-6 random field of views at 10x magnification from three sections 500 µm apart were assessed per colon. Scoring of colons from non-DSS treated mice was carried out as described by Erben et al 2014 (81), Scoring scheme 2 for colonic inflammation mediated by luminal antigens. Briefly, colon sections were scored (1-4) according to two criteria, a) the severity of inflammatory cell infiltrate and, b) epithelial changes including hyperplasia, goblet cell and crypt loss. The scores for each criteria were averaged to give the final score, with 4 representing the highest score. Scoring of colons from DSS treated mice was carried out as described by Erben et al 2014 (81). Briefly, colon sections were scored on four criteria, 1, Inflammatory cell infiltrate, 2, erosion and ulceration, 3, submucosal edema and hyperplasia and 4, loss of crypts and goblet cells, for a total clinical score of 12. Images were captured using a Leica DM 4000 or Leica DM 6000 Power Mosaic microscope (Leica Microsystems).

### Flow cytometry

Flow cytometric staining was performed as described (15). Fluorochrome-conjugated antibody clones against the following surface antigens were: CD3 (17A2), CD45.2 (104), B220 (RA3-6B2), IgD (11-26), Ly6G (IA8) (Biolegend). CD4 (RM4-5), CD8 (53-6.7), CD11b (M1/70), CD23 (B3B4), CD44 (IM7), FOXP3 (FJK-165) (Ebioscience). CD21 (7G6), CD25 (PC61.5), CD62L (MEL-14) CD138 (281-2), FAS (Jo2), IgM (II/41) (BD Biosciences). Data were acquired with FACSCanto II and FORTESSA flow cytometers (BD Biosciences) and analyzed using FlowJo software (Tree Star).

### Immunoblot analysis and immunoprecipitation

PBMC’s and murine thymocytes were lysed using lysis buffer (Cell Signaling Technology). Protein concentration was measured using the Bradford assay (Bio-Rad) and total protein (20-25 µg) resolved on a 7 – 10% SDS PAGE gel and then transferred to a nitrocellulose membrane, Immobilon-P^®^ (Merck Millipore). Membranes were incubated with anti-IκBα (product #: 9242), anti-A20 (Product #: 5630; clone: D13H3) (Cell Signaling Technology), anti-A20 (product #: ab13597; clone: 59A426) (Abcam) or anti-beta-actin (Product #: A1978; clone: AC15) (Sigma-Aldrich); followed by horseradish peroxidase (HRP)-labeled secondary antibody goat-anti-mouse IgG Fc (Product #: 31433) (Pierce Antibodies) or donkey-anti-rabbit IgG (Product #: NA934V) (GE Life Sciences). HRP conjugates bound to antigen were detected and visualized by using an ECL detection kit (GE Life Sciences).

### Real Time quantitative PCR

Total RNA was extracted using the RNeasy Plus Mini Kit (Qiagen) and reverse transcribed using Quantitect Reverse Transcription Kit (Qiagen). Primers were designed using Primer3 (82) with sequences obtained from GenBank and synthesized by Sigma Aldrich (Supplementary Table 1). PCR reactions were performed on the LightCycler^®^ 480 Real Time PCR System (Roche) using the FastStart SYBR Green Master Mix (Roche). *RPL13A* and ACTB were used as housekeeping genes and data analysed using the 2ΔΔCT method. Initial denaturation was =performed at 95° C for 10 sec, followed by a three-step cycle consisting of 95° C for 15 sec =(4.8° C/s, denaturation), 63° C for 30 sec (2.5° C/sec, annealing), and 72° C for 30 sec (4.8° C/s, elongation). A melt-curve was performed after finalization of 45 cycles at 95° C for 2 min, 40° C for 3 min and gradual increase to 95° C with 25 acquisitions/° C.

### Statistical methods

Results are expressed as mean +/- standard error mean (SEM). Statistical analysis was performed using the Student’s *t*-test or ANOVA where indicated.

## Supplementary Figures and legends

**Supplementary Figure 1.**
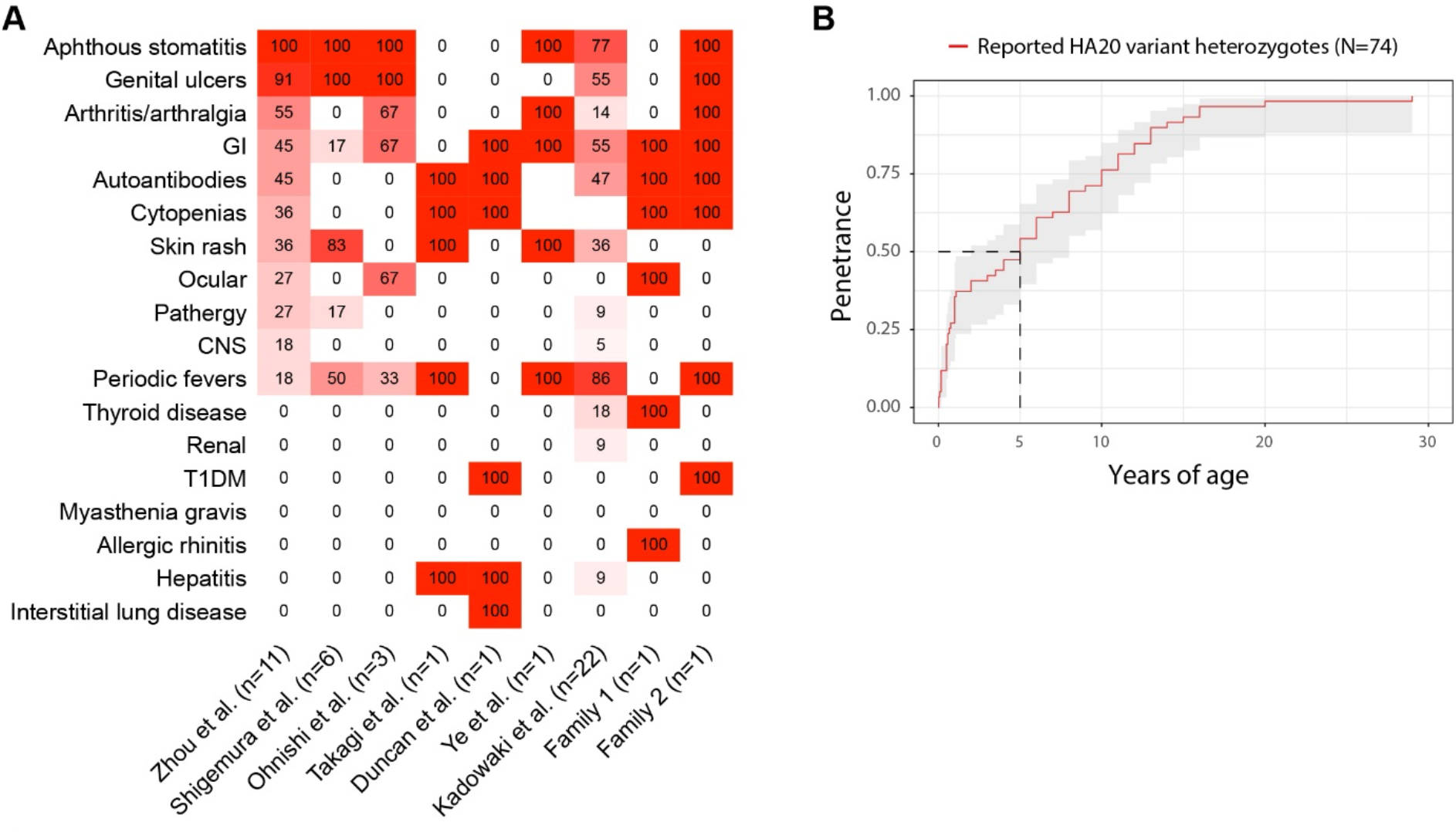
**(A)** Summary of clinical presentations observed in probands from family 1 and 2 in this study, compared to reported cases (18-22). **(B)** Age of disease of 74 reported HA20 variant heterozygotes.

**Supplementary Figure 2.**
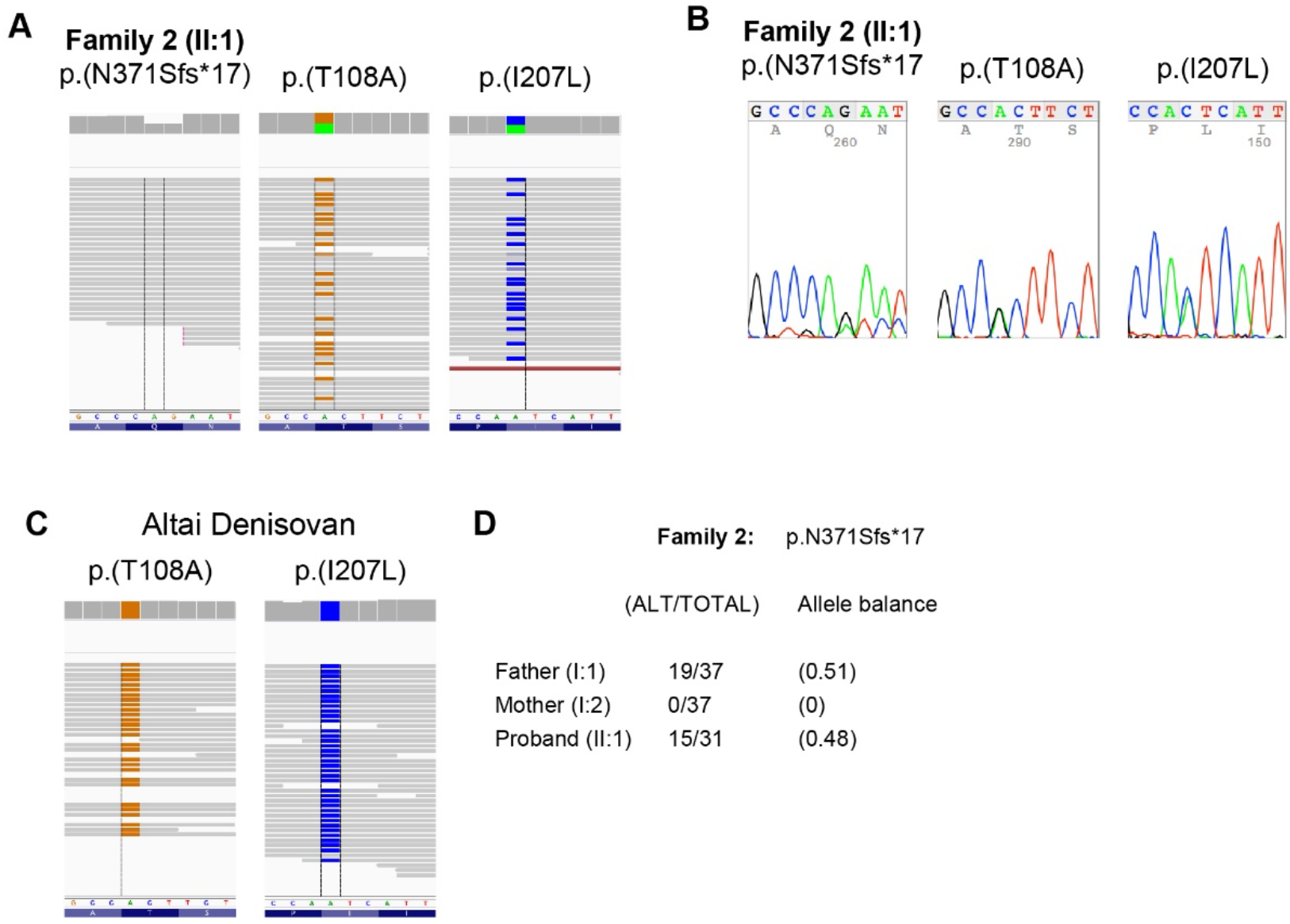
**(A)** Read data of family 2 across TNFAIP3 frame shift (N371Sfs*17) and codols 108 and 207 **(B)** and representative chromatograms. **(C)** Read data of a high-coverage Denisovan genome (83) across TNFAIP3 codons 108 and 207. **(D)** Read call outs of p.N371Sfs*17 in family 2 trio.

**Supplementary Figure 3.**
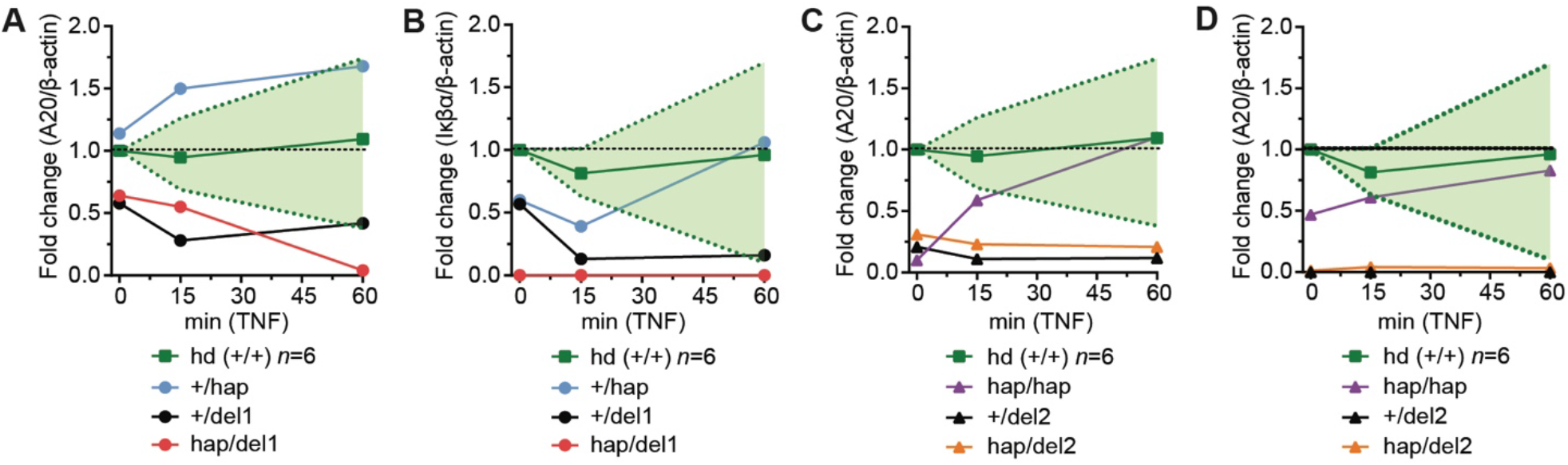
**(A-D)** Densitometry analysis of A20 (A, C) or IκBα levels (B, D) from Trio 1 (A, B) and Trio 2 (C, D) immunoblots in Fig 2A, B. Six healthy donor (Hd) values presented as +/- range (light green shade).

**Supplementary Figure 4.**
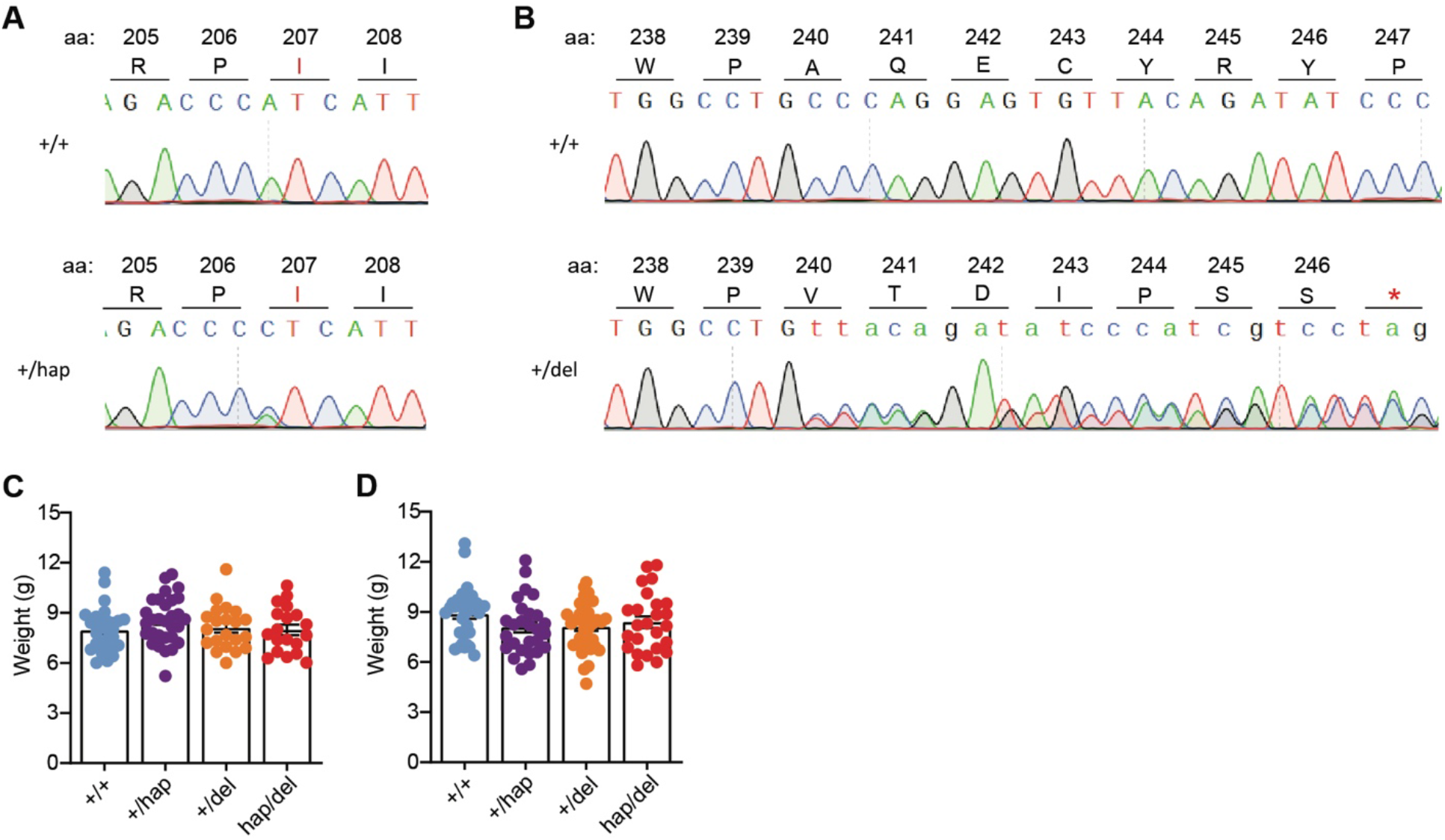
**(A)** Representative Sanger chromatogram for mice with the I207L (hap) or **(B)** A240Vfs*7 (del) alleles. **(C-D)** Weight of 8-week-old female (C) or male (D) *Tnfaip3* hap/del mice of the indicated genotype.

**Supplementary Figure 5.**
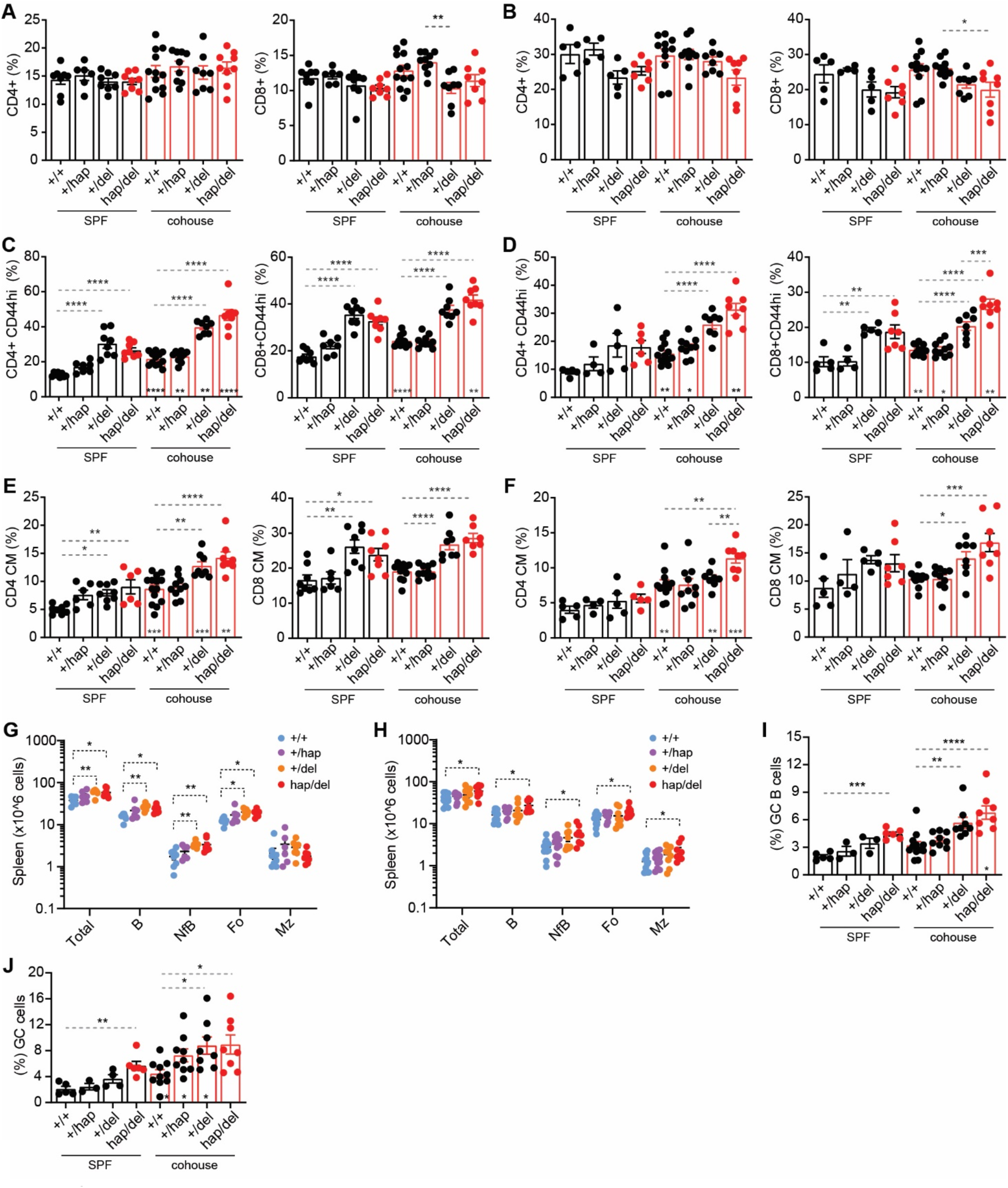
**(A-F)** Flow cytometry analysis of splenic (A, C, E) or mesenteric lymph node (B, D, F) T cell populations from Tnfaip3I207L/246tr mice housed under SPF conditions (black bars) or cohoused with dirty mice for 8 weeks (red bars). **(A, B)** Percentage of CD4+ or CD8+ T lymphocytes. **(C, D)** Percentage of CD4+ or CD8+ CD44hi memory T cells. **(E, F)** CD44hiCD62hi central memory CD8+ or CD4+ T cells. **(G, H)** Absolut numbers of B cells in the spleen of individual mice with the indicated *Tnfaip3* genotype, in SPF conditions (G) or post cohouse with dirty mice for 8 weeks (H). B, B cells; NFB, newly formed B cells; FO, follicular B cells; MZ, marginal zone B cells. **(I-J)** Flow cytometry analysis of splenic (I) or mesenteric lymph node (J) B cell germinal centres. **(K-M)** Percent proportions of splenic inflammatory monocytes (K), neutrophils (L) and NK1.1 (M) positive cells were also assessed.**n**Statistical analysis was performed by one-way anova ANOVA with Tukey post hoc test, and data shown as the mean +/- s.e.m: \**P <* 0.05; \*\**P <* 0.01; \*\*\**P <* 0.001; \*\*\*\**P <* 0.0001. ns = not significant.

**Supplementary Figure 6.**
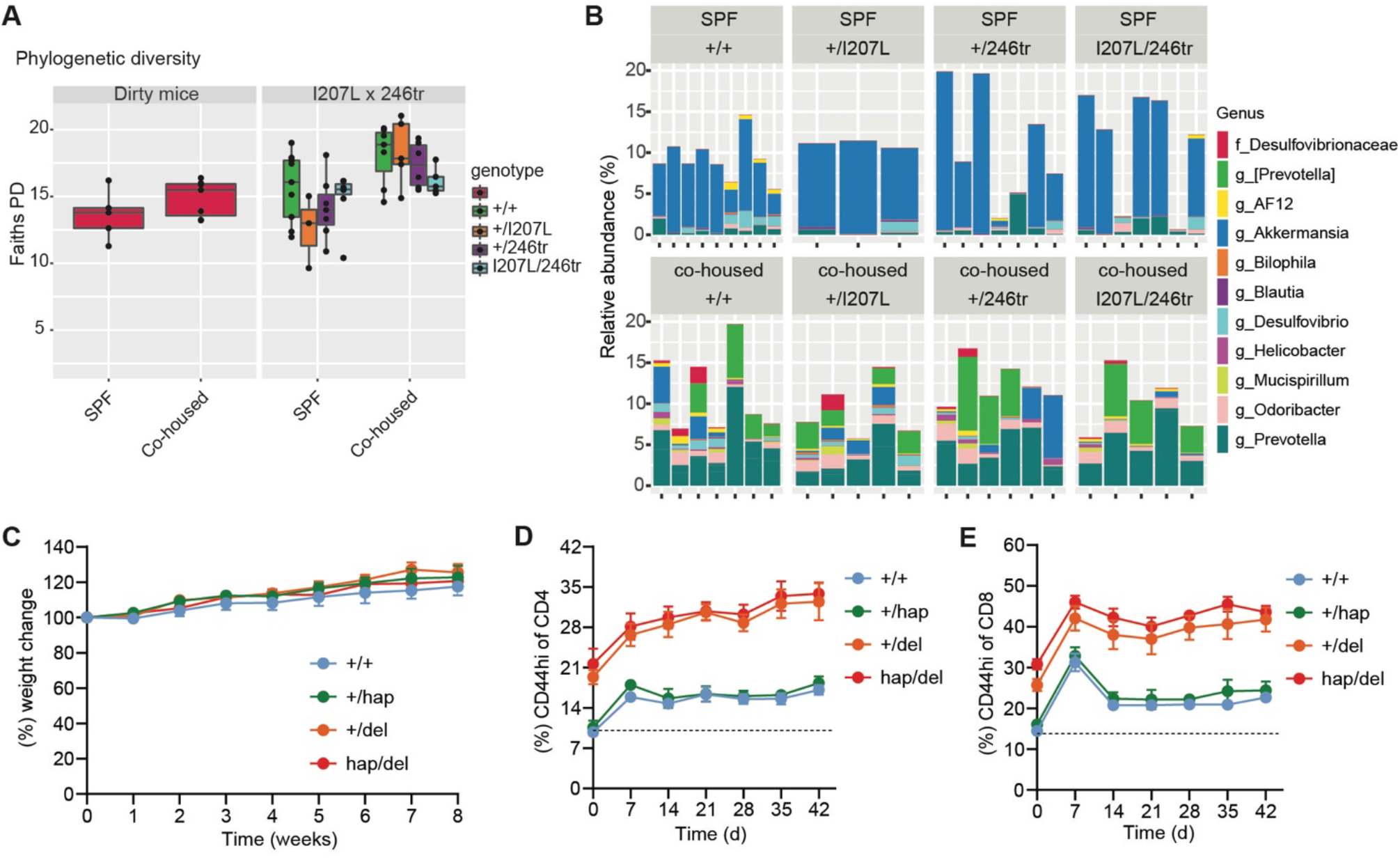
**(A)** Bar plot showing relative abundance of significantly changing bacteria species following cohousing (p<0.05, two-way ANOVA). **(B)** Weekly weights presented as percent change in starting weight, for *Tnfaip3* hap/del mice of the indicated genotypes cohoused with dirty mice (+/+ n=4; +/hap n=5; +/del n=3; hap/del n=3). **(C)** Percent CD44hi CD8+ T cells or **(D)** CD44hi CD4+ T cells in the blood of *Tnfaip3* hap/del mice co-housed with dirty mice (+/+ n=7; +/hap n=6; +/del n=5; hap/del n=5).

**Supplementary Figure 7.**
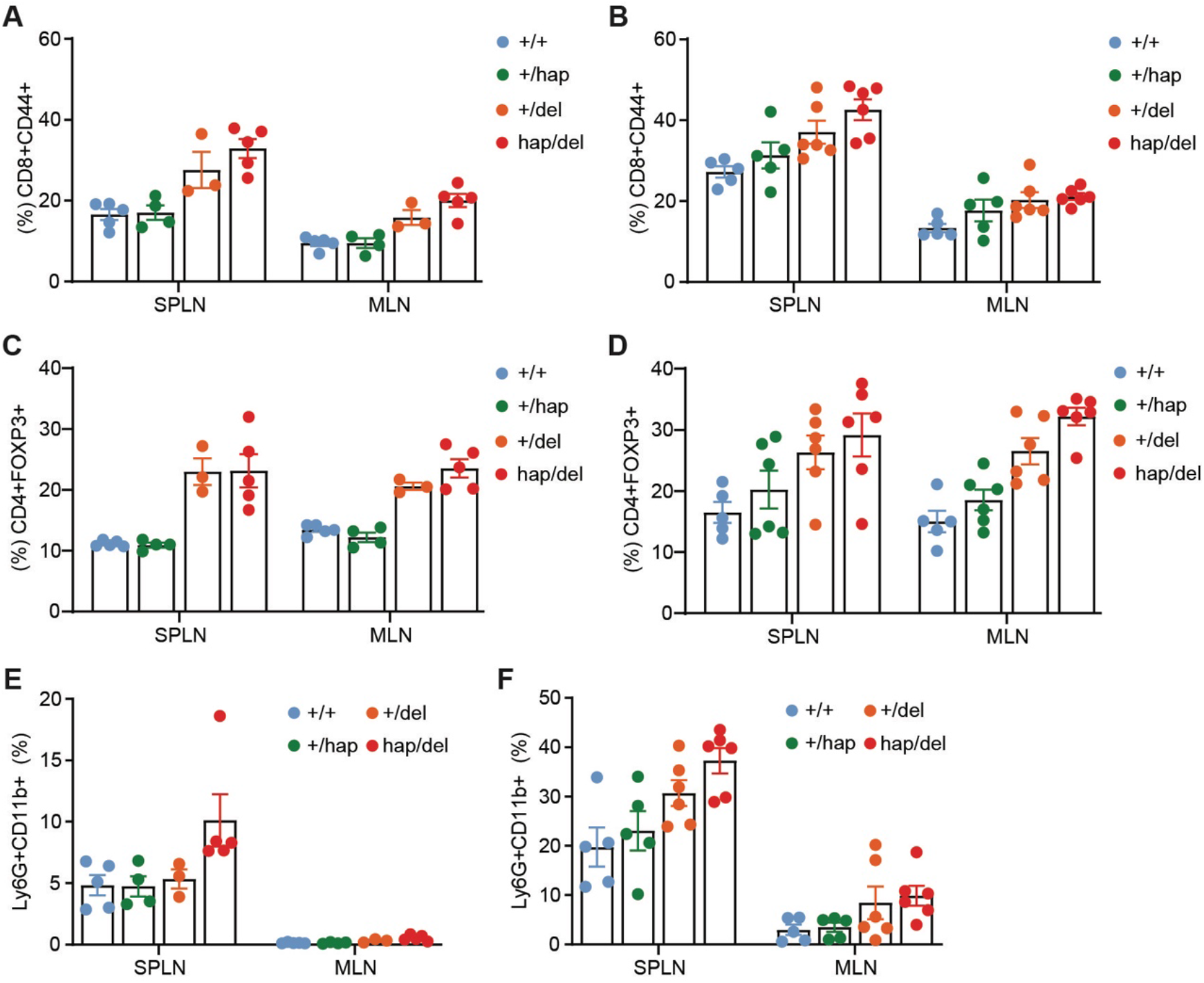
**(A, B)** Percentage of CD44hi CD8+ or CD4+ T cells from the spleen (SPLN) or Mesenteric lymph node (MLN) of *Tnfaip3* hap/del mice of the indicated genotype (B) before or (C) 14 days post DSS treatment. **(C, D)** Percentage of CD4+FOXP3+ cells from the spleen (SPLN) or Mesenteric lymph node (MLN) of *Tnfaip3* hap/del mice of the indicated genotype (D) before or (E) 14 days post DSS treatment. **(E, F)** Percentage of CD45+Ly6G+CD11b+ neutrophils from the spleen (SPLN) or Mesenteric lymph node (MLN) of *Tnfaip3* hap/del mice of the indicated genotype (F) before or (G) 14 days post DSS treatment.

## Supplementary Tables

**Supplementary Table 1.**
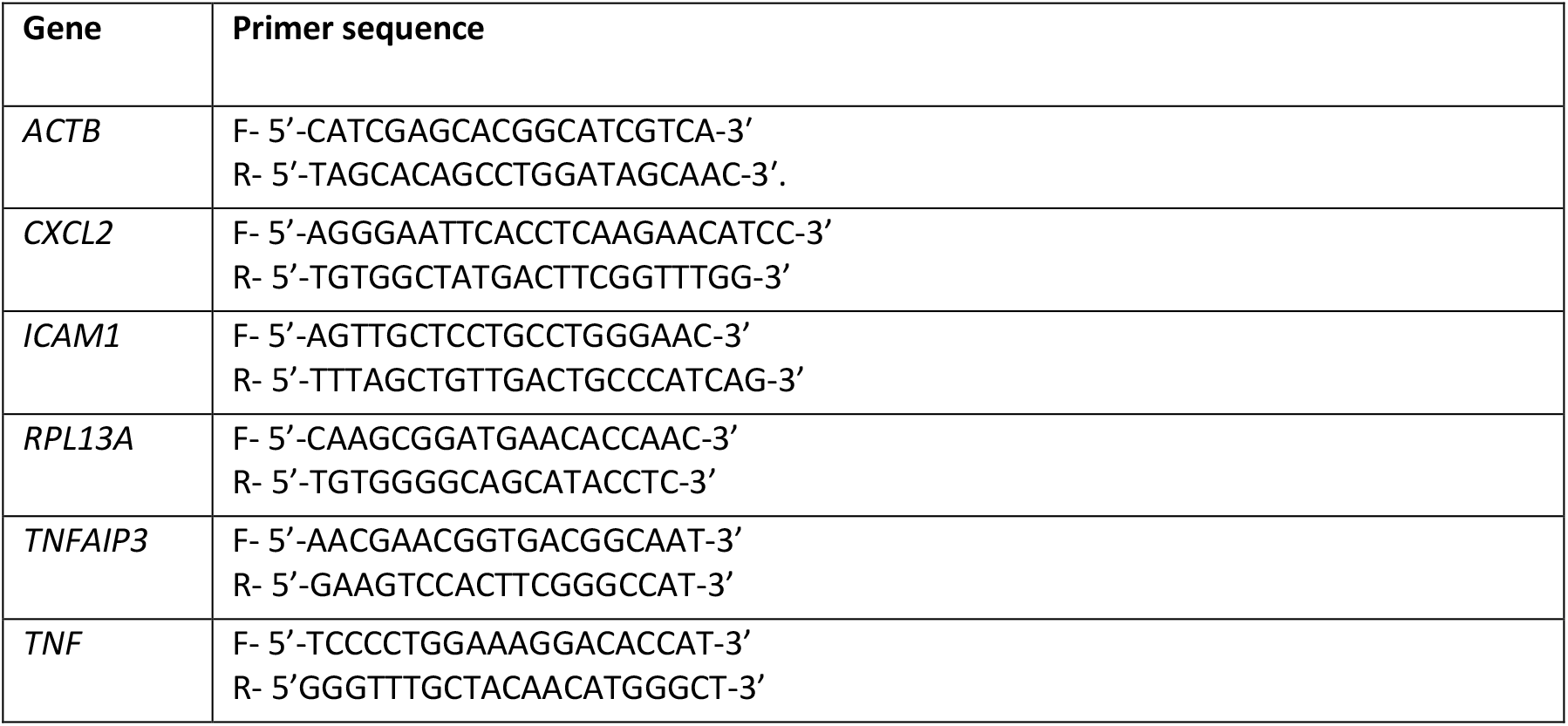
Human primers used for qRT-PCR analysis.

## REFERENCES

1. Lee EG, Boone DL, Chai S, Libby SL, et al. Failure to regulate TNF-induced NF-kappaB and cell death responses in A20-deficient mice. Science. 2000;289(5488):2350–4.

2. Boone DL, Turer EE, Lee EG, Ahmad RC, et al. The ubiquitin-modifying enzyme A20 is required for termination of Toll-like receptor responses. Nat Immunol. 2004;5(10):1052–60.

3. Cultrone D, Zammit NW, Self E, Postert B, et al. A zebrafish functional genomics model to investigate the role of human A20 variants in vivo. Scientific reports. 2020;10(1):19085.

4. Zammit NW, Walters SN, Seeberger KL, O’Connell PJ, et al. A20 as an immune tolerance factor can determine islet transplant outcomes. JCI Insight. 2019;4(21).

5. Wertz IE, O’Rourke KM, Zhou H, Eby M, et al. De-ubiquitination and ubiquitin ligase domains of A20 downregulate NF-kappaB signalling. Nature. 2004;430(7000):694–9.

6. Evans PC, Ovaa H, Hamon M, Kilshaw PJ, et al. Zinc-finger protein A20, a regulator of inflammation and cell survival, has de-ubiquitinating activity. Biochem J. 2004;378(Pt 3):727–34.

7. Lu TT, Onizawa M, Hammer GE, Turer EE, et al. Dimerization and ubiquitin mediated recruitment of a20, a complex deubiquitinating enzyme. Immunity. 2013;38(5):896–905.

8. Bosanac I, Wertz IE, Pan B, Yu C, et al. Ubiquitin binding to A20 ZnF4 is required for modulation of NF-kappaB signaling. Mol Cell. 2010;40(4):548–57.

9. Wertz IE, Newton K, Seshasayee D, Kusam S, et al. Phosphorylation and linear ubiquitin direct A20 inhibition of inflammation. Nature. 2015;528(7582):370–5.

10. Verhelst K, Carpentier I, Kreike M, Meloni L, et al. A20 inhibits LUBAC-mediated NF-kappaB activation by binding linear polyubiquitin chains via its zinc finger 7. EMBO J. 2012;31(19):3845–55.

11. Liuwantara D, Elliot M, Smith MW, Yam AO, et al. Nuclear factor-kappaB regulates beta-cell death: a critical role for A20 in beta-cell protection. Diabetes. 2006;55(9):2491–501.

12. Dixit VM, Green S, Sarma V, Holzman LB, et al. Tumor necrosis factor-alpha induction of novel gene products in human endothelial cells including a macrophage-specific chemotaxin. Journal of Biological Chemistry. 1990;265(5):2973–8.

13. Krikos A, Laherty CD, Dixit VM. Transcriptional activation of the tumor necrosis factor alpha-inducible zinc finger protein, A20, is mediated by kappa B elements. J Biol Chem. 1992;267:17971–6.

14. Hutti JE, Turk BE, Asara JM, Ma A, et al. IkappaB kinase beta phosphorylates the K63 deubiquitinase A20 to cause feedback inhibition of the NF-kappaB pathway. Mol Cell Biol. 2007;27(21):7451–61.

15. Zammit NW, Siggs OM, Gray PE, Horikawa K, et al. Denisovan, modern human and mouse TNFAIP3 alleles tune A20 phosphorylation and immunity. Nat Immunol. 2019;20(10):1299–310.

16. Ma A, Malynn BA. A20: linking a complex regulator of ubiquitylation to immunity and human disease. Nat Rev Immunol. 2012;12(11):774–85.

17. Zhou Q, Wang H, Schwartz DM, Stoffels M, et al. Loss-of-function mutations in TNFAIP3 leading to A20 haploinsufficiency cause an early-onset autoinflammatory disease. Nat Genet. 2016;48(1):67–73.

18. Duncan CJA, Dinnigan E, Theobald R, Grainger A, et al. Early-onset autoimmune disease due to a heterozygous loss-of-function mutation in TNFAIP3 (A20). Ann Rheum Dis. 2017.

19. Kadowaki T, Ohnishi H, Kawamoto N, Hori T, et al. Haploinsufficiency of A20 causes autoinflammatory and autoimmune disorders. J Allergy Clin Immunol. 2017.

20. Ohnishi H, Kawamoto N, Seishima M, Ohara O, et al. A Japanese family case with juvenile onset Behcet’s disease caused by TNFAIP3 mutation. Allergol Int. 2017;66(1):146–8.

21. Shigemura T, Kaneko N, Kobayashi N, Kobayashi K, et al. Novel heterozygous C243Y A20/TNFAIP3 gene mutation is responsible for chronic inflammation in autosomal-dominant Behcet’s disease. RMD Open. 2016;2(1):e000223.

22. Takagi M, Ogata S, Ueno H, Yoshida K, et al. Haploinsufficiency of TNFAIP3 (A20) by germline mutation is involved in autoimmune lymphoproliferative syndrome. J Allergy Clin Immunol. 2017;139(6):1914–22.

23. Aeschlimann FA, Batu ED, Canna SW, Go E, et al. A20 haploinsufficiency (HA20): clinical phenotypes and disease course of patients with a newly recognised NF-kB-mediated autoinflammatory disease. Ann Rheum Dis. 2018;77(5):728–35.

24. Berteau F, Rouviere B, Nau A, Le Berre R, et al. ’A20 haploinsufficiency (HA20): clinical phenotypes and disease course of patients with a newly recognised NF-kB-mediated autoinflammatory disease’. Ann Rheum Dis. 2019;78(5):e35.

25. He T, Huang Y, Luo Y, Xia Y, et al. Haploinsufficiency of A20 Due to Novel Mutations in TNFAIP3. J Clin Immunol. 2020;40(5):741–51.

26. Guggino WB, Stanton BA. New insights into cystic fibrosis: molecular switches that regulate CFTR. Nature reviews Molecular cell biology. 2006;7(6):426–36.

27. Wexler NS, Lorimer J, Porter J, Gomez F, et al. Venezuelan kindreds reveal that genetic and environmental factors modulate Huntington’s disease age of onset. Proc Natl Acad Sci U S A. 2004;101(10):3498–503.

28. Karczewski KJ, Francioli LC, Tiao G, Cummings BB, et al. The mutational constraint spectrum quantified from variation in 141,456 humans. Nature. 2020;581(7809):434–43.

29. Gittelman RM, Schraiber JG, Vernot B, Mikacenic C, et al. Archaic Hominin Admixture Facilitated Adaptation to Out-of-Africa Environments. Curr Biol. 2016;26(24):3375–82.

30. Karin M, Ben-Neriah Y. Phosphorylation meets ubiquitination: the control of NF-[kappa]B activity. Annu Rev Immunol. 2000;18:621–63.

31. Mathes E, O’Dea EL, Hoffmann A, Ghosh G. NF-kappaB dictates the degradation pathway of IkappaBalpha. EMBO J. 2008;27(9):1357–67.

32. Aeschlimann FA, Laxer RM. Haploinsufficiency of A20 and other paediatric inflammatory disorders with mucosal involvement. Curr Opin Rheumatol. 2018;30(5):506–13.

33. Turer EE, Tavares RM, Mortier E, Hitotsumatsu O, et al. Homeostatic MyD88-dependent signals cause lethal inflammation in the absence of A20. J Exp Med. 2008;205(2):451–64.

34. Beura LK, Hamilton SE, Bi K, Schenkel JM, et al. Normalizing the environment recapitulates adult human immune traits in laboratory mice. Nature. 2016;532(7600):512–6.

35. Rosshart SP, Herz J, Vassallo BG, Hunter A, et al. Laboratory mice born to wild mice have natural microbiota and model human immune responses. Science. 2019;365(6452).

36. Albers CA, Paul DS, Schulze H, Freson K, et al. Compound inheritance of a low-frequency regulatory SNP and a rare null mutation in exon-junction complex subunit RBM8A causes TAR syndrome. Nat Genet. 2012;44(4):435-9, S1-2.

37. Hebbel RP. Sickle hemoglobin instability: a mechanism for malarial protection. Redox Rep. 2003;8(5):238–40.

38. Phornphutkul C, Introne WJ, Perry MB, Bernardini I, et al. Natural history of alkaptonuria. N Engl J Med. 2002;347(26):2111–21.

39. Blau N, van Spronsen FJ, Levy HL. Phenylketonuria. Lancet. 2010;376(9750):1417–27.

40. Hakansson A, Tormo-Badia N, Baridi A, Xu J, et al. Immunological alteration and changes of gut microbiota after dextran sulfate sodium (DSS) administration in mice. Clin Exp Med. 2015;15(1):107–20.

41. Johansson ME, Gustafsson JK, Sjoberg KE, Petersson J, et al. Bacteria penetrate the inner mucus layer before inflammation in the dextran sulfate colitis model. PLoS One. 2010;5(8):e12238.

42. Johansson ME, Gustafsson JK, Holmen-Larsson J, Jabbar KS, et al. Bacteria penetrate the normally impenetrable inner colon mucus layer in both murine colitis models and patients with ulcerative colitis. Gut. 2014;63(2):281–91.

43. Vereecke L, Sze M, Mc Guire C, Rogiers B, et al. Enterocyte-specific A20 deficiency sensitizes to tumor necrosis factor-induced toxicity and experimental colitis. J Exp Med. 2010;207(7):1513–23.

44. Scher JU, Sczesnak A, Longman RS, Segata N, et al. Expansion of intestinal Prevotella copri correlates with enhanced susceptibility to arthritis. Elife. 2013;2:e01202.

45. Iljazovic A, Roy U, Galvez EJC, Lesker TR, et al. Perturbation of the gut microbiome by Prevotella spp. enhances host susceptibility to mucosal inflammation. Mucosal Immunol. 2020.

46. Atarashi K, Tanoue T, Ando M, Kamada N, et al. Th17 Cell Induction by Adhesion of Microbes to Intestinal Epithelial Cells. Cell. 2015;163(2):367–80.

47. Larsen JM. The immune response to Prevotella bacteria in chronic inflammatory disease. Immunology. 2017;151(4):363–74.

48. Caruso R, Mathes T, Martens EC, Kamada N, et al. A specific gene-microbe interaction drives the development of Crohn’s disease-like colitis in mice. Sci Immunol. 2019;4(34).

49. Kullberg MC, Ward JM, Gorelick PL, Caspar P, et al. Helicobacter hepaticus triggers colitis in specific-pathogen-free interleukin-10 (IL-10)-deficient mice through an IL-12-and gamma interferon-dependent mechanism. Infect Immun. 1998;66(11):5157–66.

50. Gobert AP, Sagrestani G, Delmas E, Wilson KT, et al. The human intestinal microbiota of constipated-predominant irritable bowel syndrome patients exhibits anti-inflammatory properties. Scientific reports. 2016;6:39399.

51. Zhai R, Xue X, Zhang L, Yang X, et al. Strain-Specific Anti-inflammatory Properties of Two Akkermansia muciniphila Strains on Chronic Colitis in Mice. Front Cell Infect Microbiol. 2019;9:239.

52. Vereecke L, Vieira-Silva S, Billiet T, van Es JH, et al. A20 controls intestinal homeostasis through cell-specific activities. Nature communications. 2014;5:5103.

53. Altin JA, Tian L, Liston A, Bertram EM, et al. Decreased T-cell receptor signaling through CARD11 differentially compromises forkhead box protein 3-positive regulatory versus T(H)2 effector cells to cause allergy. J Allergy Clin Immunol. 2011;127(5):1277–85 e5.

54. Ronin E, Lubrano di Ricco M, Vallion R, Divoux J, et al. The NF-kappaB RelA Transcription Factor Is Critical for Regulatory T Cell Activation and Stability. Frontiers in immunology. 2019;10:2487.

55. Oh H, Grinberg-Bleyer Y, Liao W, Maloney D, et al. An NF-kappaB Transcription-Factor-Dependent Lineage-Specific Transcriptional Program Promotes Regulatory T Cell Identity and Function. Immunity. 2017;47(3):450–65 e5.

56. Zhang Z, Zhang W, Guo J, Gu Q, et al. Activation and Functional Specialization of Regulatory T Cells Lead to the Generation of Foxp3 Instability. J Immunol. 2017;198(7):2612–25.

57. Mikami N, Kawakami R, Chen KY, Sugimoto A, et al. Epigenetic conversion of conventional T cells into regulatory T cells by CD28 signal deprivation. Proc Natl Acad Sci U S A. 2020;117(22):12258–68.

58. Salama AD, Chitnis T, Imitola J, Ansari MJ, et al. Critical role of the programmed death-1 (PD-1) pathway in regulation of experimental autoimmune encephalomyelitis. J Exp Med. 2003;198(1):71–8.

59. Free ME, Bunch DO, McGregor JA, Jones BE, et al. Patients with antineutrophil cytoplasmic antibody-associated vasculitis have defective Treg cell function exacerbated by the presence of a suppression-resistant effector cell population. Arthritis Rheum. 2013;65(7):1922–33.

60. Valencia X, Yarboro C, Illei G, Lipsky PE. Deficient CD4+CD25high T regulatory cell function in patients with active systemic lupus erythematosus. J Immunol. 2007;178(4):2579–88.

61. Ooi JD, Snelgrove SL, Engel DR, Hochheiser K, et al. Endogenous foxp3(+) T-regulatory cells suppress anti-glomerular basement membrane nephritis. Kidney Int. 2011;79(9):977–86.

62. Wang KW, Zhan X, McAlpine W, Zhang Z, et al. Enhanced susceptibility to chemically induced colitis caused by excessive endosomal TLR signaling in LRBA-deficient mice. Proc Natl Acad Sci U S A. 2019;116(23):11380–9.

63. Chen H, Li H, Liu Z. Interplay of intestinal microbiota and mucosal immunity in inflammatory bowel disease: a relationship of frenemies. Therap Adv Gastroenterol. 2020;13:1756284820935188.

64. Mevissen TE, Kulathu Y, Mulder MP, Geurink PP, et al. Molecular basis of Lys11-polyubiquitin specificity in the deubiquitinase Cezanne. Nature. 2016;538(7625):402–5.

65. Siggs OM, Russell A, Singh-Grewal D, Wong M, et al. Preponderance of CTLA4 Variation Associated With Autosomal Dominant Immune Dysregulation in the MYPPPY Motif. Frontiers in immunology. 2019;10:1544.

66. Gayevskiy V, Roscioli T, Dinger ME, Cowley MJ. Seave: a comprehensive web platform for storing and interrogating human genomic variation. bioRxiv. 2018.

67. Minoche AE, Lundie B, Peters GB, Ohnesorg T, et al. ClinSV: Clinical grade structural and copy number variant detection from whole genome sequencing data. medRxiv. 2020(2020.06.30.20143453).

68. Bolyen E, Rideout JR, Dillon MR, Bokulich NA, et al. Reproducible, interactive, scalable and extensible microbiome data science using QIIME 2. Nature biotechnology. 2019;37(8):852–7.

69. Martin M. Cutadapt removes adapter sequences from high-throughput sequencing reads. EMBnetjournal. 2011;17(1):10–2.

70. Rognes T, Flouri T, Nichols B, Quince C, et al. VSEARCH: a versatile open source tool for metagenomics. PeerJ. 2016;4:e2584.

71. Amir A, McDonald D, Navas-Molina JA, Kopylova E, et al. Deblur Rapidly Resolves Single-Nucleotide Community Sequence Patterns. mSystems. 2017;2(2).

72. Bokulich NA, Kaehler BD, Rideout JR, Dillon M, et al. Optimizing taxonomic classification of marker-gene amplicon sequences with QIIME 2’s q2-feature-classifier plugin. Microbiome. 2018;6(1):90.

73. Mandal S, Van Treuren W, White RA, Eggesbo M, et al. Analysis of composition of microbiomes: a novel method for studying microbial composition. Microb Ecol Health Dis. 2015;26:27663.

74. McMurdie PJ, Holmes S. phyloseq: an R package for reproducible interactive analysis and graphics of microbiome census data. PLoS One. 2013;8(4):e61217.

75. Hamady M, Lozupone C, Knight R. Fast UniFrac: facilitating high-throughput phylogenetic analyses of microbial communities including analysis of pyrosequencing and PhyloChip data. ISME J. 2010;4(1):17–27.

76. M.J. A. Permutational multivariate analysis of variance (PERMANOVA). In: N. Balakrishnan TC, B. Everitt, W. Piegorsch, F. Ruggeri and J.L. Teugels, editor. Wiley StatsRef: Statistics References Online 2017.

77. Oksanen JF, Blanchet G, Friendly M, Kindt R, et al. Vegan: Community Ecology Package. R package version 2.5-6. https://CRAN.R-project.org/package=vegan2019n

78. Faith DP. Conservation evaluation and phylogenetic diversity. Biological Conservation. 1992;61(1):1–10.

79. Kembel SW, Cowan PD, Helmus MR, Cornwell WK, et al. Picante: R tools for integrating phylogenies and ecology. Bioinformatics. 2010;26(11):1463–4.

80. Wickham H., François R, Henry L, Müller K. dplyr: A Grammar of Data Manipulation. R package version 1.0.2. https://CRAN.R-project.org/package=dplyr2020

81. Erben U, Loddenkemper C, Doerfel K, Spieckermann S, et al. A guide to histomorphological evaluation of intestinal inflammation in mouse models. Int J Clin Exp Pathol. 2014;7(8):4557–76.

82. Untergasser A, Cutcutache I, Koressaar T, Ye J, et al. Primer3--new capabilities and interfaces. Nucleic Acids Res. 2012;40(15):e115.

83. Meyer M, Kircher M, Gansauge MT, Li H, et al. A high-coverage genome sequence from an archaic Denisovan individual. Science. 2012;338(6104):222–6.

